# Schema-induced shifts in mice navigational strategies are unveiled by a minimal behavioral model of spatial exploration

**DOI:** 10.1101/2020.12.21.423808

**Authors:** Christina-Anna Vallianatou, Alejandra Alonso, Adrian Aleman, Lisa Genzel, Federico Stella

## Abstract

Shifts in spatial patterns produced during the execution of a navigational task can be used to track the effects of the accumulation of knowledge and the acquisition of structured information about the environment. Here we provide a quantitative analysis of mice behavior while performing a novel goal localization task in a large, modular arena, the HexMaze. To demonstrate the effects of different forms of previous knowledge we first obtain a precise statistical characterization of animals’ paths with sub-trial resolution and over different phases of learning. The emergence of a flexible representation of the task is accompanied by a progressive improvement of performance, mediated by multiple, multiplexed time scales. We then use a generative mathematical model of the animal behavior to isolate the specific contributions to the final navigational strategy. We find that animal behavior can be accurately reproduced by the combined effect of a goal-oriented component, becoming stronger with the progression of learning, and of a random walk component, producing choices unrelated to the task and only partially weakened in time.

## Introduction

The problem of learning and especially of the integration of new information into an already existing knowledge structure is at the center of the effort to understand brain functioning (Alonso et al., 2020). When using rodent animal models, such problem has often been addressed in the context of spatial navigation and map learning (Richards et al., 2014; Wang and Morris, 2010). Animals knowledge about the environment and the degree to which they can acquire new information can be linked to their ability to easily navigate to specific locations and flexibly adapt to changes in the environment (Behrens et al., 2018). Nevertheless, the characterization of the effects of learning has been mostly restricted to simple tasks with limited spatial and temporal complexity, focusing on isolating specific components of the learning process with highly-controlled paradigms. Indeed, the difficulties in precisely monitoring the animal behavior pose one of the major limiting factors in the development of more comprehensive experimental paradigms (Fonio et al., 2009). Here we aim at filling this gap by providing a quantitative framework for the description of navigational strategies expressed by mice while completing a goal-reaching task.

Assessing the effects of the accumulation of learning on the performance in a spatial orientation task requires the combination of two elements. On the one hand the complexity of the task should be high enough to allow for the expression of rich behavioral patterns and of different grades of information acquisition (Benjamini et al., 2011). Disentangling the different components informing animal choices requires providing animals with multiple options over a sizable spatial and temporal interval. Such availability is also a requirement in the interest of understanding animal behavior in its naturalistic setting (Tchernichovski and Benjamini, 1998). Wild rodents experience will include an articulate system of burrows together with the surrounding layout, a situation that can only be captured in the laboratory by studying spatial learning in larger, more complex environments (Wood et al., 2018).

As a consequence of the richer behavioral repertoire accessible to the animal, successfully tracking the evolution of task-related abilities requires the deployment of specific quantification tools, aimed not only at measuring task performance but also the specifics of animal behavior that accompany it (Dvorkin et al., 2008). Such tools should also provide a link between observable changes in the animal choice patterns and shifting navigational strategies underlying such choices (Gehring et al., 2015; Ruediger et al., 2012).

In this study we use a novel navigational task, featuring a goal-localization paradigm with extended spatial and temporal dimensions, the HexMaze (Figure 1, top). Mice learn to locate a reward location in a larger, modularly structured maze providing precise control over animals’ paths. Testing animals over a long temporal period, and after a modification of the environment as either we introduce a novel reward location or alternatively we place a set of barriers to block some paths (Figure 1), we look at the effects of previous knowledge on their performance and on their ability to flexibly incorporate novel information. We are thus able to track different contributions to the observed animal behavior, including those linked with information encoding, memory consolidation and schema acquisition (for a complete description of learning dynamics on the maze see (Alonso et al., n.d.)). To test the effects of such components we apply statistical analysis to obtain a fully-characterized picture describing the evolution of animal choices with sub-trial resolution. Importantly, such detailed phenomenological description of trial-by-trial behavioral profiles is then complemented with a generative model of task trajectories based on a mathematical description of task completion process (Figure 1).

**Figure 1:**
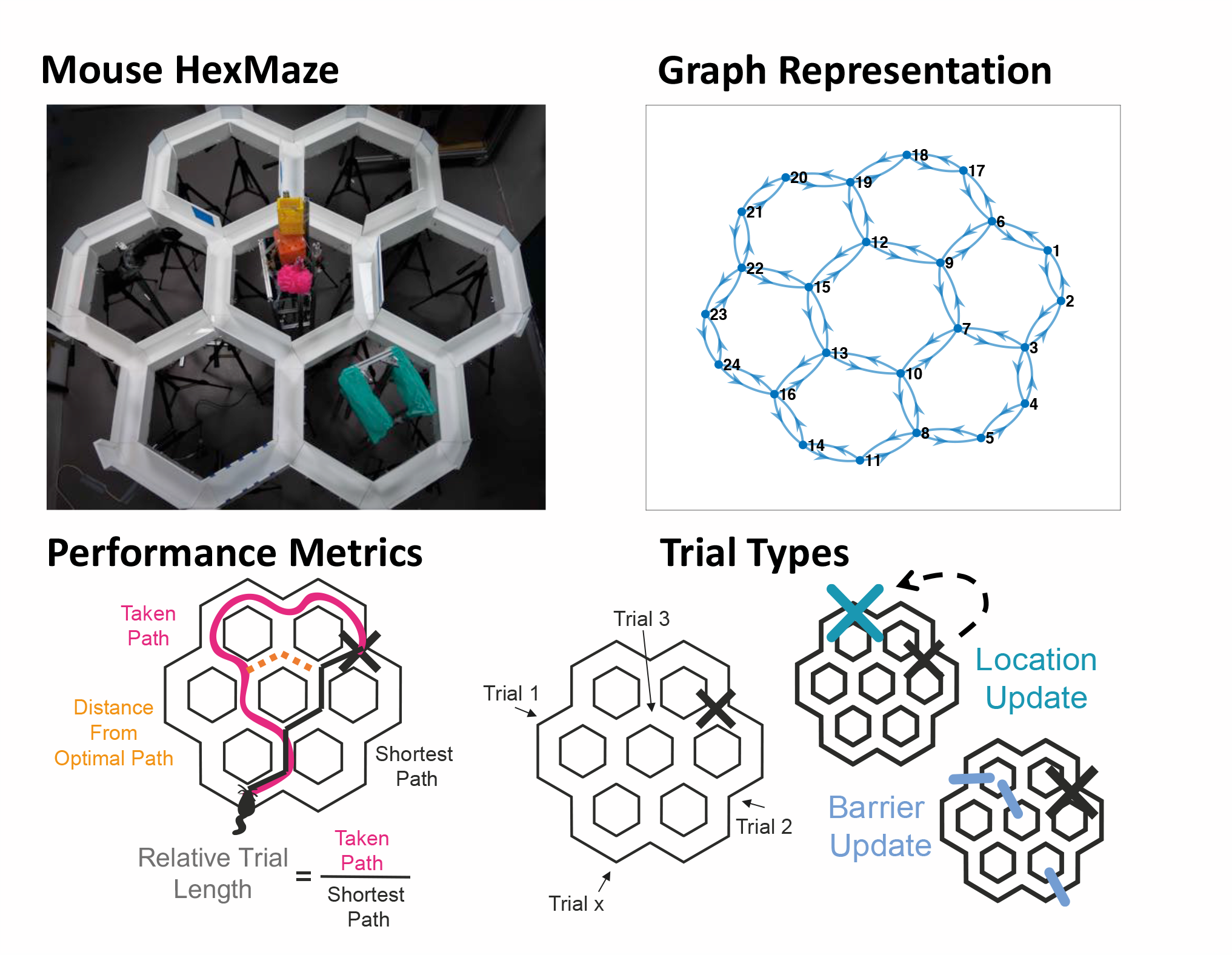
HexMaze Structure and Experimental Paradigm. **Top**: View of the maze (left) and its graph representation used in the analysis (right). **Bottom Left**: Two main performance metrics are used. 1) Relative Trial Length is the length of the paths taken by the animal divided by the shortest possible path to the goal location. 2) Distance from Optimal Path is the distance of the animal position at any time from the closest point of the shortest path. **Bottom right:** During training, animals started each trial from a different location and had to navigate to a fixed Goal Location. After the animals had acquired the general maze knowledge during the Build-Up, Updates were performed with inclusion of new barriers (Barrier Update) or new goal locations (Location Update)

This modeling approach unveils a limited set of principles guiding animal choices. Our results show how animal behavior can be faithfully reproduced by a minimal mixture of random walking and goal-directed runs over limited distances. The relative importance of these two components over the different phases of learning (initial goal-location acquisition, consolidation over multiple sessions, schema update in coincidence with environmental modifications) not only provides a concise characterization of animal approach to the task, but mirrors the emergence of task-specific memory constructs, offering a direct quantification of learning induced patterns of behavior. We find that, although we can observe an increasing amount of knowledge about the task and the maze structure being incorporated by animals, their performance never really converges to be completely goal-oriented. Instead we can measure the influence of a consistent random component, interfering with optimal task performance, and being only partially reduced with increasing familiarity to the task. Persistence of such task-independent activity could be a product of exposing animals to an expanded task complexity, effectively giving them increased freedom to divert from task completion, but it might also reflect mice specific idiosyncratic behavior, possibly triggered by the specie propensity to hyperactivity (Jones et al., 2017). In both cases, further application of our modeling approach is likely to provide further insight on the diversity of behavioral approaches linked to different cognitive demands and different species.

## Results

### The HexMaze Experiment

The HexMaze is arranged as six regular densely packed hexagons, forming twelve two-way and twelve three-way choice points (nodes) 36.3 cm apart, in total spanning 2 m × 1.9 m (Fig. 1, Top). Gangways between nodes were 10 cm wide and flanked by either 7.5 cm or 15 cm tall walls. Maze floor and walls were white and opaque, with local and global cues applied in and around the maze to enable easy spatial differentiation and good spatial orientation; overall leading to a complex, integrated maze. During training food was placed in one of the nodes and the animal had to learn to navigate efficiently from different start locations to the goal location (GL).

Animals went through two phases of training: Build-Up and Updates. In the Build-Up the animals should create a cognitive map of the maze environment; in contrast, during Updates, stable performance is achieved and they should be simply updating the cognitive map. These two phases also differed in the frequency of GL switches: during Build-Up, the GL remained stable for five and more sessions, while during Updates a change occurred every three sessions (see also below). Different Update types were performed: including barriers in the environment (Barrier Update) and changing the goal location (Location Update) (Fig. 1, bottom).

### Characterization of animal behavior in the HexMaze

We first set out to quantify the time-evolution of animal behavior. The structure of the HexMaze allows for an efficient tracking of the animal choices, as its behavior is easily described by the sequence of visited maze nodes. Therefore, in the following we will focus on this descriptor to measure different aspects of the animal performance during the experiment. Clearly, the main element to be taken into account when analyzing behavior is the ability of the animal to efficiently localize the reward and reach it through a path as short as possible (Figure 1).

A perfectly optimal, goal-oriented behavior, would imply that after a necessary learning transient the trajectories selected by the animal would progressively converge toward the shortest available given the start location and the reward one. By measuring the ratio between the length of the actual path (measured in terms of number of nodes visited before reaching the goal) and the length of the optimal path (Relative Trial Length = RTL), over a certain amount of trials, one would then expect to observe this distribution to be more and more skewed towards the value of 1, corresponding to the animal actually following the optimal path (Figure 2). Indeed what we find is a progressive increase in the percentage of trials with a low ASR score, within each session, across sessions and also for each of the experimental conditions (Figure 2, last column). Multiple effects point to an actual presence of learning and to a growing awareness of the maze structure and goal location in the animals. First, during Build-Up not only the score improves within each session, but animals consistently do better in the first trial of a new session compared to the previous one. Then, while the score goes back to pre-learning values at the beginning of the Location Update, when a new unknown goal is introduced, it reaches its asymptotic value faster compared to Build-up. And finally, the insertion of barriers has only a very limited effect on the animal performance. At the same time the hypothesis of over-wise, totally committed mice is challenged by the fact that although increasing in time, the probability of a perfect (or almost perfect) run remains substantially below 1 over the entire arc of the experiment, even after the animals have been repeatedly exposed to the maze and to a specific reward location. The trial relative length distribution shows a long tail of values larger than 1 (Figure 2, columns 1 to 3), indicating that the animal choices are far from corresponding to a purely optimal, goal-oriented strategy.

**Figure 2:**
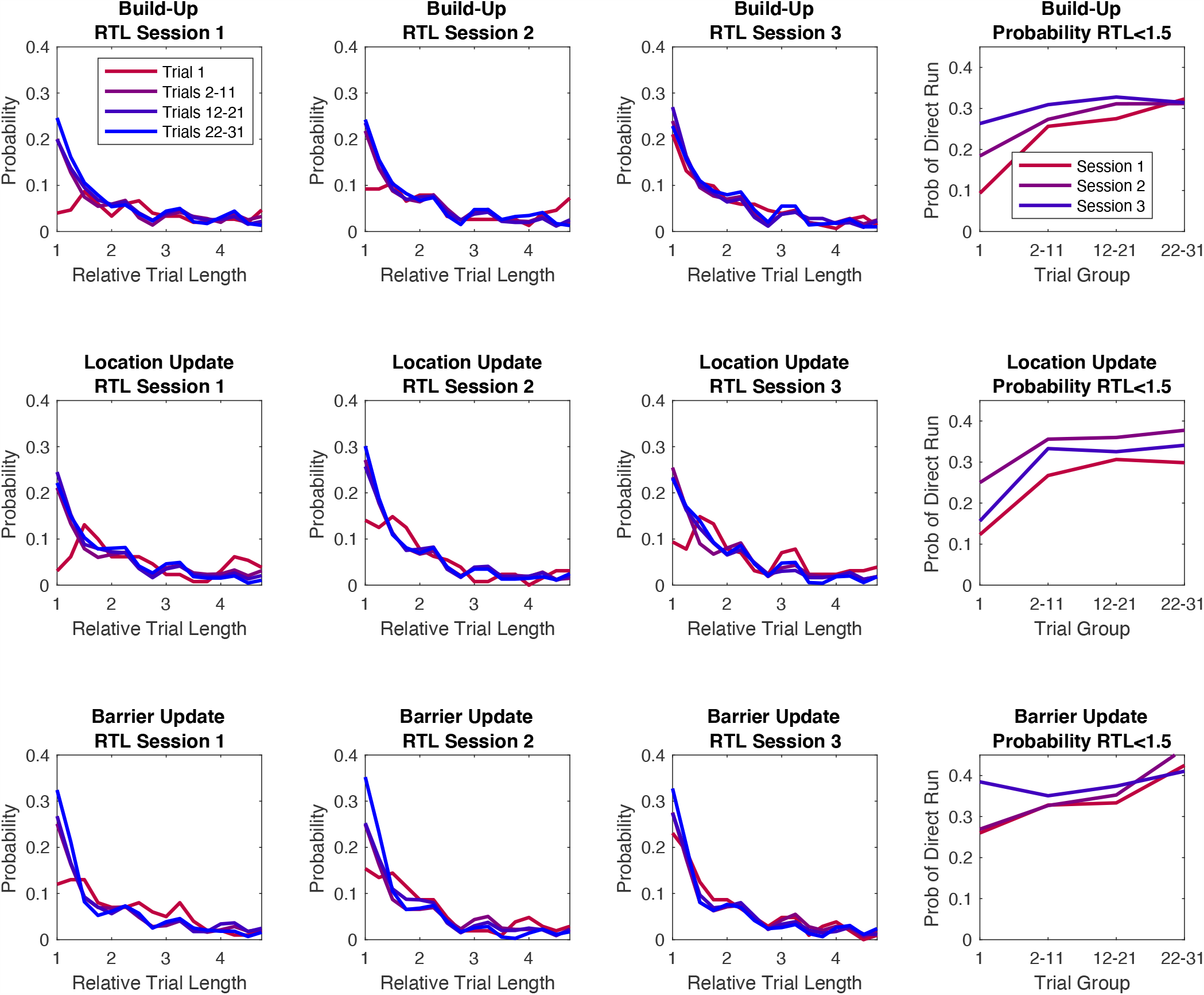
Animals are progressively more likely to take shorter paths to the goal. Distribution of relative trial lengths (RTL) for different trial groups and sessions. RTL = 1 corresponds to perfect trial. Last panel: probability of RTL < 1.5 (optimal trial) over time. For each session, trials from different animals were grouped in 4 categories: 1^st^ trial only, from 2^nd^ to 11^th^ trial, from 12^th^ to 21^st^ trial and from 22^nd^ to 31^st^ trial.

One way to further characterize the degree to which the mouse behavior is goal-directed is to measure how far its trajectory would steer away from the optimal path joining the starting location to the reward. For each trial and for each node visited by the animal during the trial, we then compute the Distance From Optimal Path (DFOP), that is, the distance between the visited node and any of the nodes comprising the optimal path. For each trial we then obtain a measure of ‘stray’ over time, providing a profile of the animal approach to the goal (Figure 3). When averaged over different set of trials, this profile shows a bump shape, quantifying the amount of deviation from optimal behavior, showing an increase at the beginning of the trial and eventually converging again towards the correct path. These curves provide us with different information about the animal trajectories over the course of the experiment: i) the amount of ‘stray’, that is the average maximum distance from the optimal path, ii) the average length of the trials and iii) how fast the animal will go back to the correct path after straying away in the beginning. We can extract this information from the data by fitting a parametrized function to the DFOP average profiles. We use a normalized difference of Gaussians (see Methods) that provides us an excellent approximation of the experimental curve shapes (Figure 4, top row). This fit depends on 3 parameters: 1. The maximum height; 2. The amount of steps to reach the maximum and 3. The amount of steps to go back to the optimal path. Plotting the value of these three parameters obtained by fitting the data from different stages of learning (Figure 4), we observe how all of them show a gradual decrease with time, consistently with the improving performance of the animal. Again, also these parameters show an overall trend across all the phases of the experiment although their evolution is not monotonous, but rather has a seesaw shape due to the partial rollbacks happening between the last trials of one session and the first trial of the following one. Together with the previous analysis, our quantification of the animal behavior, shows how the navigation of mice in the HexMaze, can be described as a combination of learning-based choices (evident in the progressive improvement in all goal-related metrics) and of a persistent non-optimal component, keeping the overall behavior away from perfect performance even for late sessions and trials.

**Figure 3:**
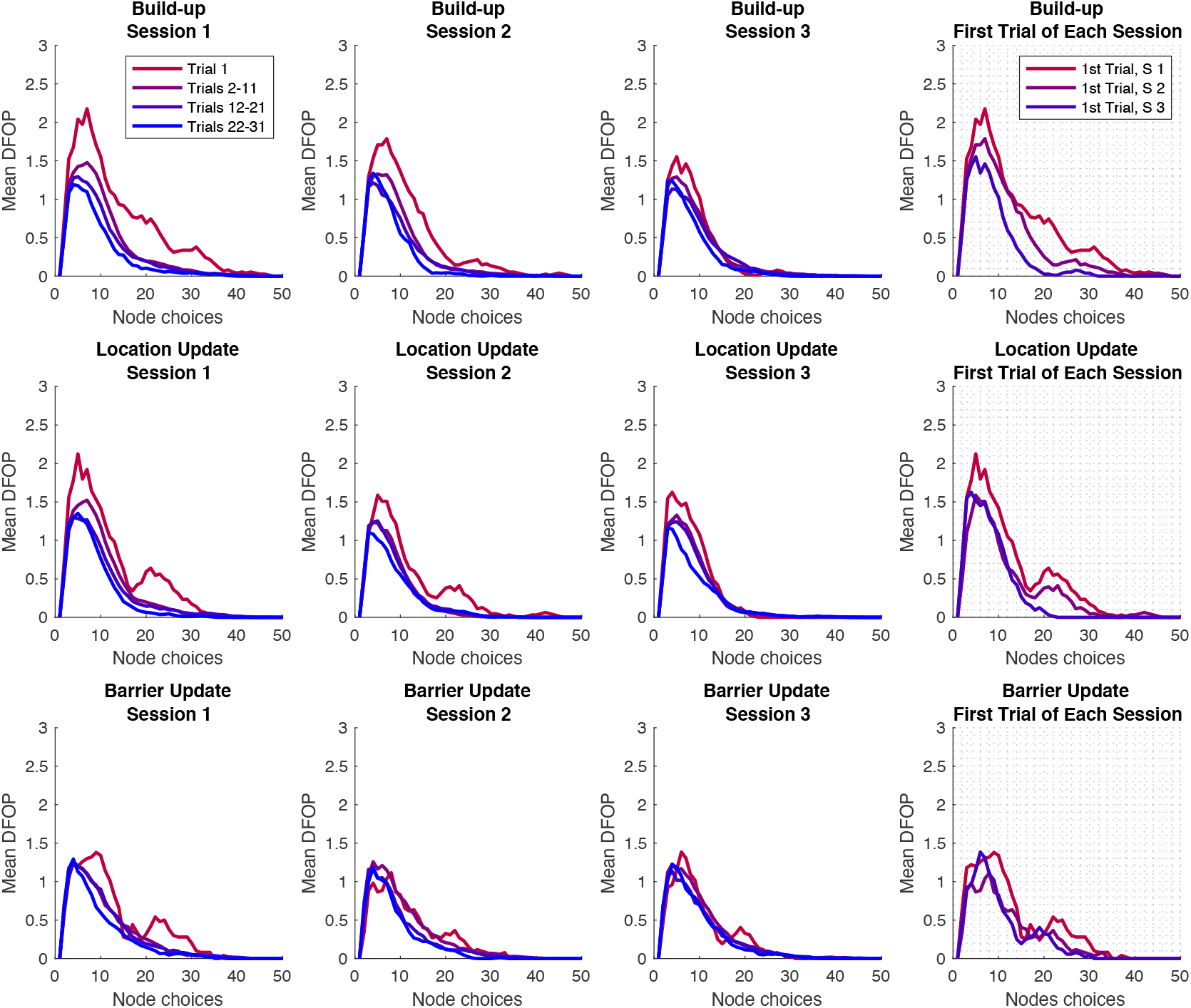
Quantification of animal trajectories departure from optimal path. Distance from optimal path (DFOP) over time for all trial groups and sessions. Last column panels: comparison of the 1st trial for the 3 sessions. The effects of learning can be seen in the progressive reduction of the distance within each session. Additionally, DFOP decreases on the first trial of every successive session. Once a new goal location is introduced, convergence to asymptotic performance is faster than during initial learning. Finally, the insertion of barriers has only limited effects on behavior.

**Figure 4:**
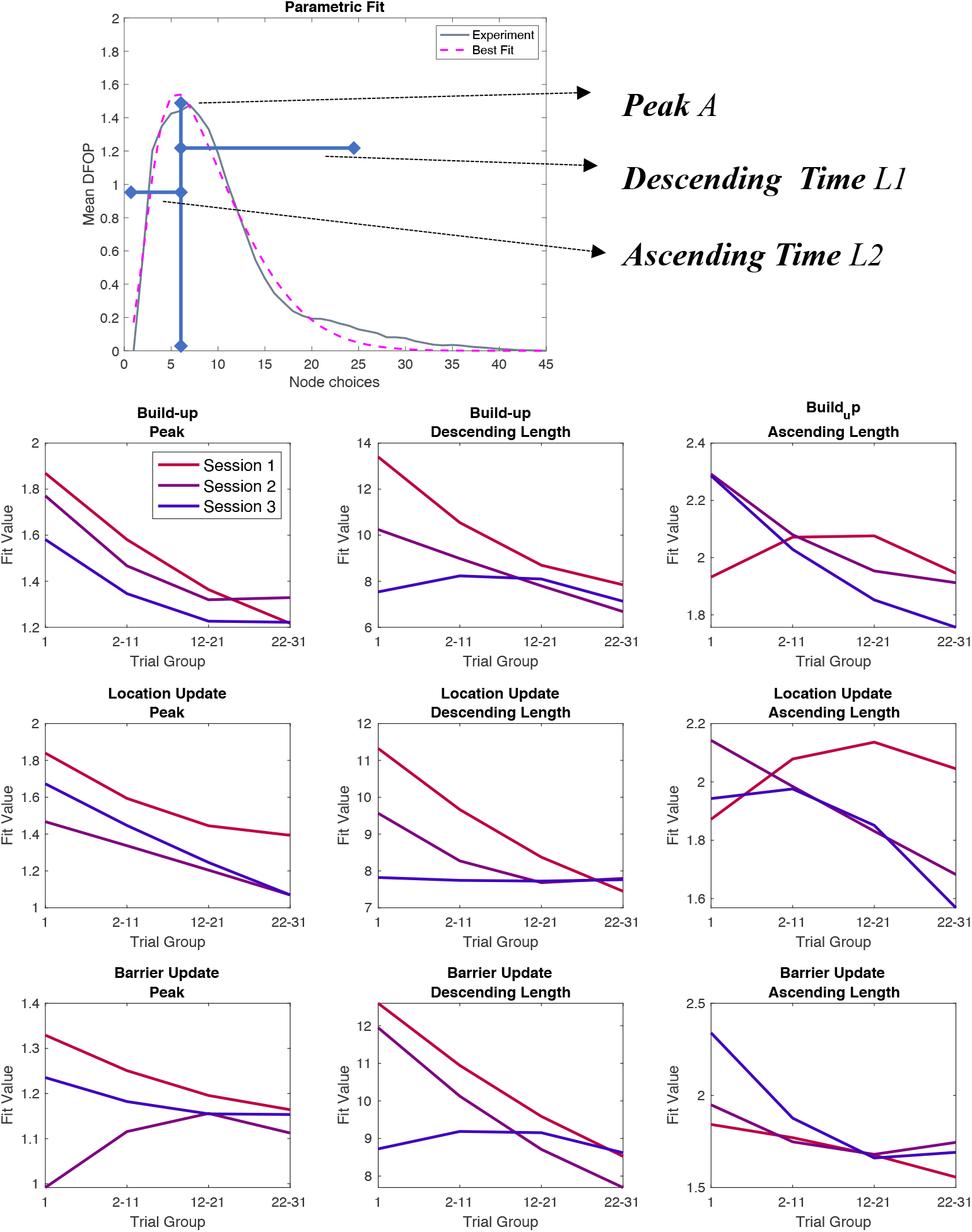
Learning sharpens animal performance by progressively reducing trajectory distance from optimal path. Results of the parametric fit of the curves in Figure 3. Top row, example of the obtained match between experimental curves and the parametric fit. Bottom, value of the fit parameters over time for all conditions and sessions. This fit allows us to quantify: i) the amount of ‘stray’, that is the average maximum distance from the optimal path, ii) the average length of the trials and iii) how fast the animal will go back to the correct path after straying away in the beginning. All these quantities show a modulation over time, consistently with learning effects within one session, across sessions, and during goal location shift.

### Minimal Mathematical Model Describing Animal Choices

We then asked whether such results could be reproduced by a simplified model of the animal behavior. We simulate the trajectories produced by a virtual agent navigating the same HexMaze used in the experiment as it searches for the reward location. In these simulations the mouse moves in the environment selecting the next node to visit according to a set of predefined rules. We define these rules as a combination of a random walk through the environment and direct goal-runs based on the knowledge of the reward location. Crucially, the model depends on only two parameters, η, the probability of taking a long diagonal run while randomly moving through the environment; and F, determining the probability that the animal will at any time start to run directly towards the reward location (a quantity that for this reason, we named Foresight). We thus simulate the animal behavior using different combinations of these 2 parameters and compare the obtained statistics with those collected during the real experiment. The comparison is based on quantifying the distance between the distribution of relative trial lengths in real and virtual trajectories.

We find that our simple behavioral models very accurately approximate the animal strategy in every part of learning. For each set of trials, we find a combination of η and F that makes the distributions not significantly different (p>0.1, as evaluated using Kolmogorov-Smirnov statistics) (Figure 5). We thus consider the (η, F) pair that minimizes the distance between experimental and simulation statistics for a specific set of trials (Figure 5). This pair of values is taken as best describing the behavioral characteristics of the animal navigation through the maze. The evolution of these values in time provides us with a measure of the effects of learning on animal’s behavior. Moreover, the model relevance is corroborated by finding that the simulated trajectories obtained with parameters optimized to fit the trial length distribution also reproduced other statistical features of the animal behavior. In fact, both the distribution of maximal distance from the optimal path, and that of average time spent on the inner vs. outer ring of the maze were captured by our simulations (Comparison between behavior and model: KS p>0.1, Figure 6).

**Figure 5:**
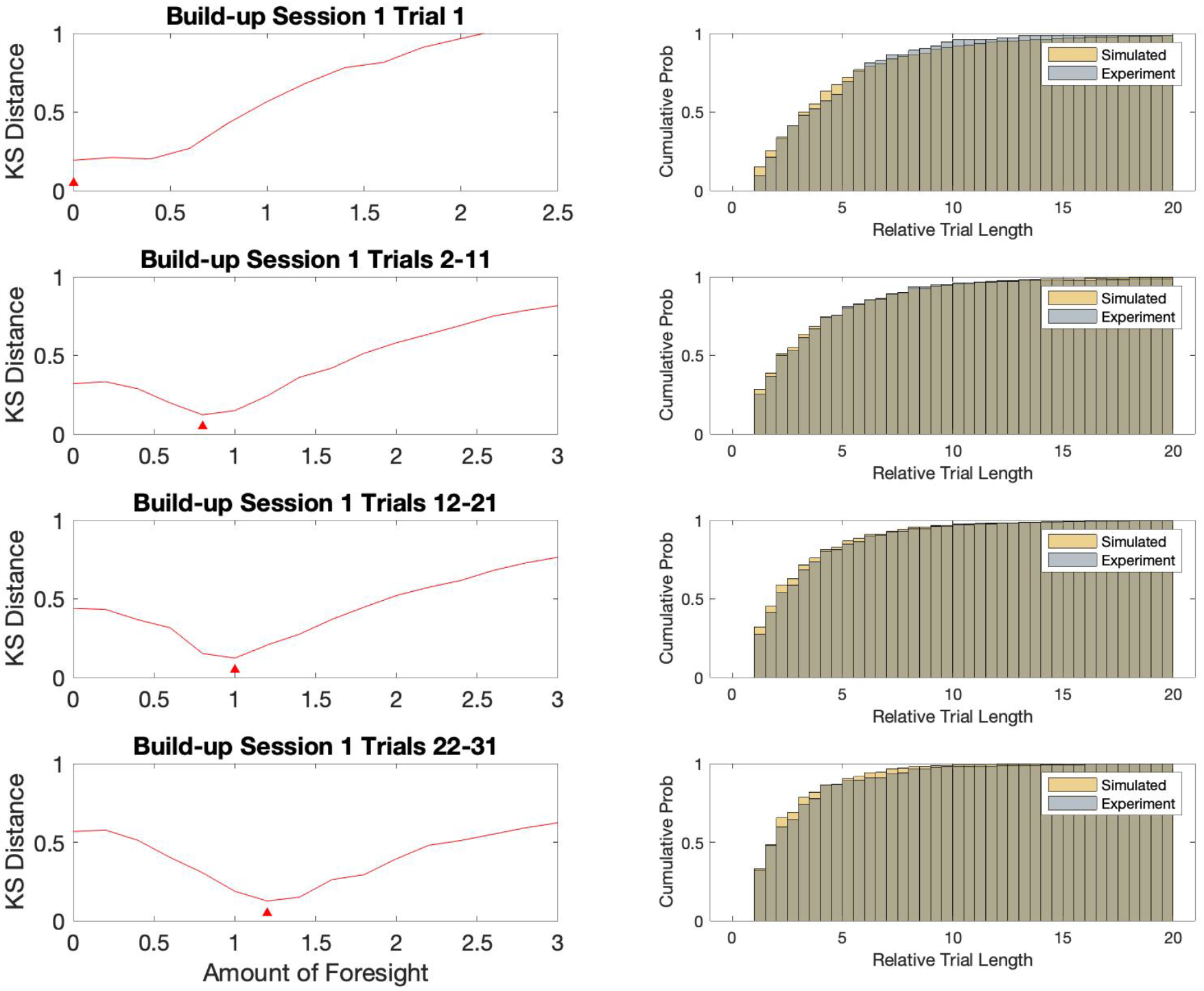
A minimal mathematical model reproduces the main properties of animal behavior. Model Fitting to Experimental Data. Left Column: Simulated-Experimental KS Distance for a range of tested Foresight values. Red triangles indicate location of best-fit F value for different sets of trials. Right Column: Comparison of cumulative distributions for different trial groups and corresponding simulation results with best F-value. All shown data is from Build-Up phase.

**Figure 6:**
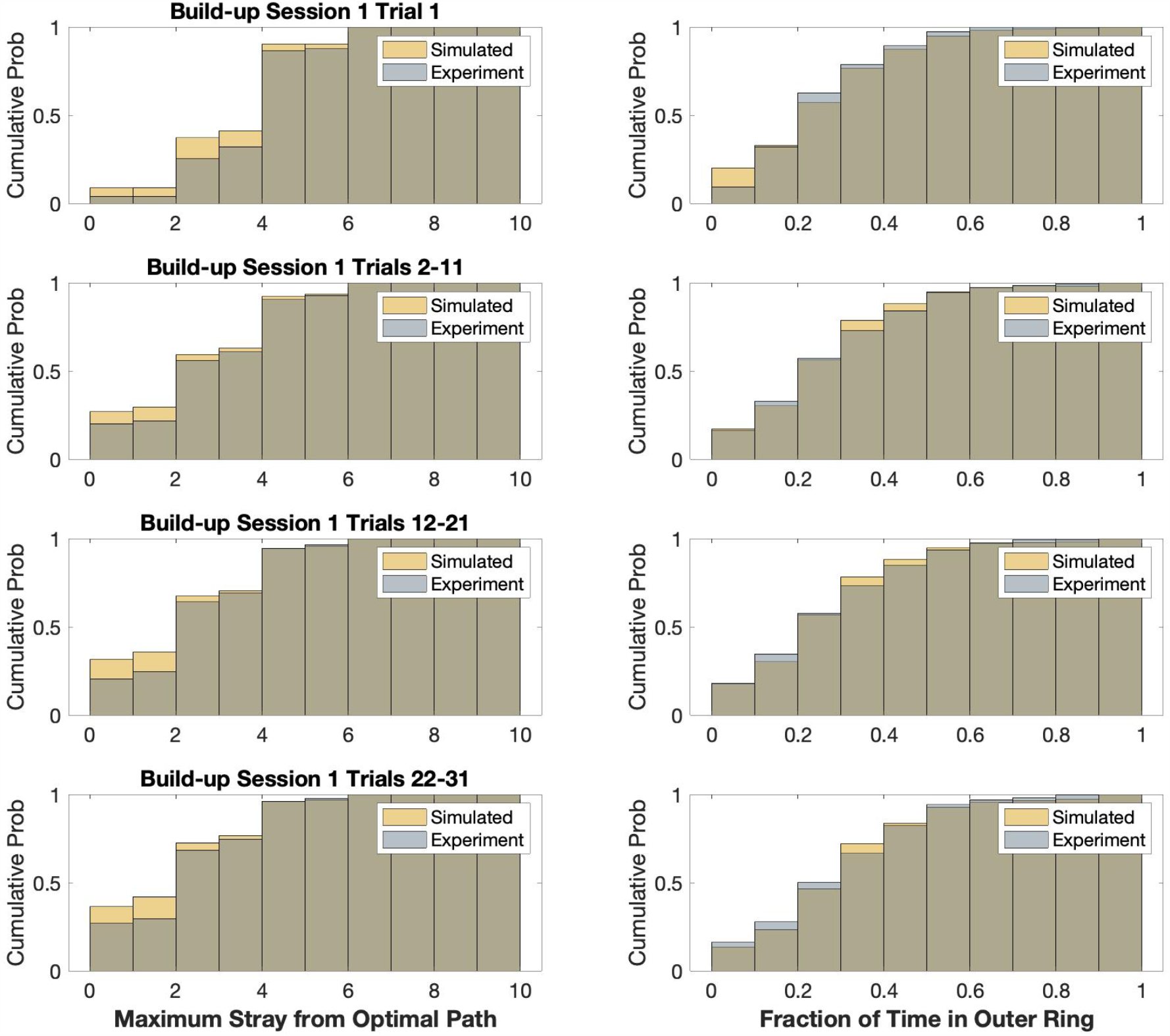
Further behavioral features matched by model fits. Same data as Figure 5. Using the same parameters obtained from fitting Relative Trial Length distributions, the model also reproduces other aspects of animal behavior such as the distribution of Maximal Distance from the Optimal Path (left) and the distribution of relative time spent in the outer or inner ring of the maze on each trial (right). Real and model-based distributions are non-significantly different for every experimental phase and trial group (KS p>0.1).

Studying the time evolution of the model parameter we first determine that the best value of η is not significantly affected by the progression of learning, and that it remains confined to 0 across the entire experiment. Therefore, learning does not change the properties of random movement across the maze, and indeed this movement pattern appears to be largely unstructured, being captured by a simple sequence of random turns. On the other hand, the effects of learning are instead reflected in the evolution of the best value for Foresight (Figure 7). As shown in the figure, its value starts at 0 for the first trial of the Build-Up phase, compatibly with an animal with no knowledge of the reward location and only randomly moving across the maze. F then progressively increases with the accumulation of trials, indicating a growing awareness for the location of the reward, its relationship to visual cues and possibly for the geometrical structure of the maze itself. Interestingly, Foresight increase is significant (Kolmogorov-Smirnov (KS) p<0.05 comparing model distributions) both across trials within one session (single lines in the plot) and across the first trial for each session, indicating a non-monotonic increase in performance (as we already found while analyzing the animal trajectories), reflecting a drop in performance between the end of one session and the start of the next one. What sort of conclusions can be drawn from the model results about the animal behavior in the maze? Even at late stages of the Build-Up phase, the Foresight value remains relatively low, never raising above a value of 2. This limit points to a significant presence of random walking even for mice that have completed a substantial number of trials. They appear to initiate goal-directed runs only when in close proximity to the reward and only rarely from the very beginning of the trial.

**Figure 7:**
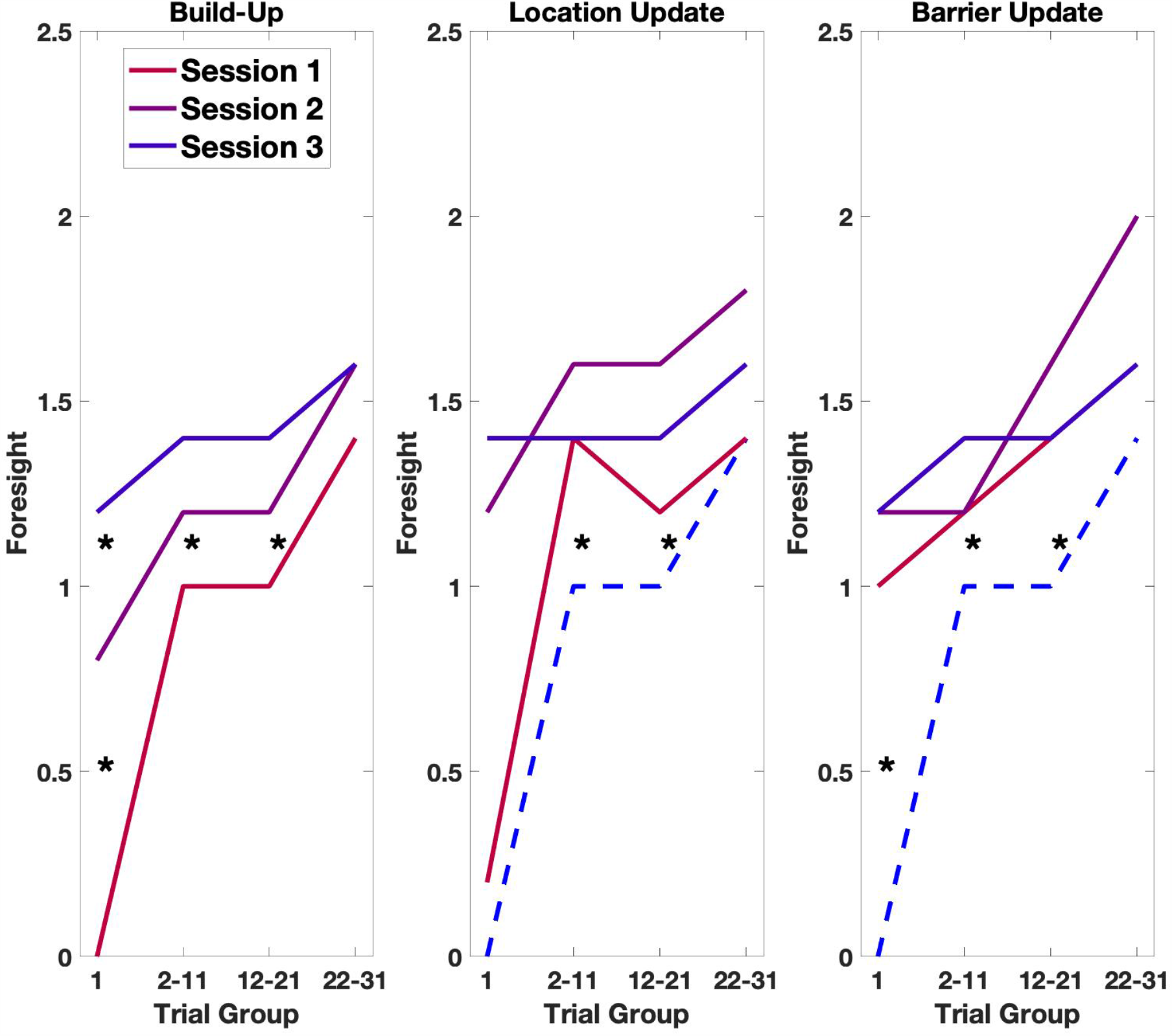
Modelling of animal behavior shows the accumulation of spatial information over the course of the experiment. Foresight Evolution: Best-fit values of Foresight for different experimental phases, sessions and trial groups. Our model reproduces the different phases of learning identified from behavioral analysis. The Foresight quantity appears to increase over the course of a session and with the accumulation of sessions. Goal location change is followed by a return to pre-learning values, but successive increase in faster than during initial task learning. Also in terms of inferred navigational strategy, the insertion of barriers in the maze has only limited effects.

Should the persistence of random behavior be taken as proof of a failure of the animal to build a complete “cognitive-map” of the maze? We can partially address this issue by looking at the performance of the model in different experimental conditions. Taking the Location Update phase, we see how Foresight is again close to 0 for the very first trial, consistently with the presence of a novel reward location. Nevertheless, in the following trials, the value of F increases at a significant faster rate compared to the build-up phase (KS p<0.05). The mice are thus able to quickly integrate the new information into the knowledge they accumulated in the previous build-up sessions. This effect is not limited to the first update session but can be seen in a rapid saturation of F in the following session, although again its value does not grow beyond 2.

Similarly, using Barrier Update sessions, when barriers are added to the maze, while maintaining the reward locations stable, the behavior of the animal appears to be at the same level of the build-up trials from the very beginning of this phase. Improvement in the goal-directed behavior can be still seen within one session, but no significant difference can be observed between sessions. The relatively low impact of barrier introduction is consistent with animal strategy being dominated by a random walk and only affected by the presence of the goal when in the proximity of it.

## Discussion

The HexMaze provides the ideal setting to characterize animal spatial cognition as it combines the availability of options to express complex behavior, with the possibility to precisely monitor and quantify animal navigational choices (Alonso et al., n.d.). In particular, in this set of experiments we leverage on its structure to study the development of goal-directed behavior in mice learning to localize a reward location. We first provide a statistical characterization of the animal navigational patterns as they traverse the maze, by comparing them to the path expected from optimal goal-oriented behavior. Our results show a clear effect of learning in mice, as they progressively tune their trajectories to reach the reward in a shorter time and visiting less nodes on the maze. We find that their trajectories are less likely to stray away from the optimal path, and that eventual detours are shorter lasting. Indeed, we can show that all of the trajectory quantifiers evolve according to a superposition of different time courses: i) they steadily improve on the first trial of each session; ii) their improvement over the course of a session becomes faster in later sessions; iii) the rate of improvement is enhanced after a novel reward location is introduced during the Update phase in contrast to the Build-Up phase; iv) introducing path-blocking barriers in the maze has only a very limited effects when the reward location is already familiar to the animal. Such sharpening of trajectories over the course of the experimental paradigm is in direct agreement with the presence of different forms of previous knowledge, all contributing to enhance the animal performance in the task (Gire et al., 2016). The emergence of an allocentric representation of the maze, the linking of specific cues to the proximity of the goal location, the strengthening of memory encoding and consolidation, can be all considered to affect the measured properties of navigational patterns.

At the same time our measures also bring to the foreground how mice behavior remains significantly distant from a purely optimal one. Mice never develop a completely goal-oriented pattern of movements, as a consistent part of their choices on the maze appears to be independent of the goal location. To identify the nature of this layer of non-optimal behavior we develop a computational model of HexMaze navigation producing virtual animal trajectories based on specific generative principles. This minimal mathematical model indicates how two components are sufficient to reproduce, with little parameter tuning, most of the statistical properties of mice real behavior. As expected from the previous results we find that the first of these components is an increasing awareness of proximity of the goal location, whose effects are felt further and further away from the goal with the accumulation of learning. Goal-directed runs stemming from this component are however combined with a basis of purely random choices that remains present throughout the experimental paradigm (Thompson et al., 2018).

One could find rather surprising this persistent neglect of the task requirements, leading to mice spending a considerable amount of time exploring portions of the maze distant from the goal, even when other behavioral measures indicate their awareness of its actual location. It is possible that such tendency could have been hidden in other experimental paradigms by the lack of options available to the animals, as they were given very little possibilities to show behavior not related to the task. In the context of a larger spatial arena the balance between “exploratory” and “exploitative” behavior (Jackson et al., 2020; Wilson et al., 2021) might shift sensibly toward the former, leading to an increase in undirected behavior. We are also aware that this pattern of behavior could be specific to mice. In fact, random exploration might be a consequence of mice hyperactivity (Jones et al., 2017) and their reluctance to consistently focus on the accomplishment of a specific task. A tendency that in the present case could be further exacerbated by the absence of sheltered locations in the maze, an element that has been shown to be of great relevance for these animals’ sense of security. With this in mind, it is not too far out to expect rats to perform very differently in this same experimental setting, which is currently under investigation. In a previous study, Jones et al (Jones et al., 2017) could show that rats and mice differed in levels of baseline activity measured as shuttle rate during inter-trial intervals; mice shuttled two to three times as frequently as rats. Species differences in behavioural ecology may underlie this difference as for example mice needing to move rapidly when outside burrows in order to minimise predation risk. They tend to use bursts of speed to run from a more sheltered position to the next.

Regardless of the final explanation of this finding, we acknowledge how these results have significant consequences for the interpretation of the neural correlates of goal-directed navigation. Place cell activity in the rodent hippocampus has been shown to organize in sequential “sweeps” linking the current location of the animal to that of one or more goal locations, when the animal is asked to take a decision (Wikenheiser and Redish, 2015). Similarly, it has been proposed that sequences of activated place cells during a Sharp Wave Ripple bear information about future navigational paths (Pfeiffer and Foster, 2013). Our results, demonstrating the co-existence of goal-oriented behavior with a substantial amount of random choices when testing mice in a more naturalistic experimental setting, suggests that such neural episodes might be circumscribed both in time and space, and might play a more limited role when considering animals with a richer set of behavioral options.

## Methods

### Subjects

Five cohorts of four male C57BL/6J mice each (Charles River Laboratories) aged two months at arrival, were group-housed in the Translational Neuroscience Unit of the Centraal Dierenlaboratorium (CDL) at Radboud University Nijmegen, Netherlands. They were kept at a 12 h light/ 12 h dark cycle and were before training food deprived overnight during the behavioural testing period. Weight was targeted to be at 90% to 85% of the animals’ estimated free-feeding weight. All animal protocols were approved by the Centrale Commissie Dierproeven (CCD, protocol number 2016-014-018). The first cohort (coh 1) was used to establish general maze and task parameters and are not included in this data set..

#### HexMaze

The HexMaze was assembled from 30 10 cm wide opaque white acrylic gangways connected by 24 equilateral triangular intersection segments, resulting in 36.3 cm distance center-to-center between intersections (Fig. 1A). Gangways were enclosed by either 7.5 cm or 15 cm tall white acrylic walls. Both local and global cues were applied to provide visual landmarks for navigation. Barriers consisted of transparent acrylic inserts tightly closing the space between walls and maze floor as well as clamped plates to prevent subjects bypassing barriers by climbing over the walls. The maze was held 70 cm above the floor to allow easy access by the experimenters.

#### Behavioural Training

After arrival and before training initiation, mice were handled in the housing room daily for 1 week (until animals freely climbed on the experimenter) and then habituated to the maze in two 1 h sessions (all four cage mates together) with intermittent handling for maze pick-ups (tubing (<u>Gouveia & Hurst</u>, <u>2017</u>)). Mice were trained either on Mondays, Wednesdays and Fridays (coh 1-3) or Tuesday and Thursday (coh 4+5). Per training day (session) each mouse underwent 30 min of training in the maze, resulting in up to 30 trials per session, Fig. S3. The maze was cleaned with 70% ethanol between animals (later clean wipes without alcohol to avoid damaging the acrylic), and to encourage returning in the next trial, a heap of food crumbles (Coco Pops, Kellogg’s, USA) was placed at a previously determined GL, which varied for each animal. GLs were counterbalanced across animals, as well as within animals across GL switches. E.g. one out of four animals, and one out of four GL per animal would be located on the inner ring of the maze while the others were on the outer ring (to shape animal behaviour against circling behaviour). Start locations for each day were generated based on their relation to the GL and previous start locations (locations did not repeat in subsequent trials, at least 60% of the trials had only one shortest path possible, first trial was different to the last and first trial of the previous session and locations had at least two choice points distance to each other as well as the GL). On average 30 start locations were needed per day per mouse, which were generated the day before training. After the mouse reached the food and ate a reward, the animal would be manually picked up with a tube, carried around the maze to disorient the mouse, and placed at the new start location. All pick-ups in the maze were done by tubing (<u>Gouveia & Hurst</u>, <u>2017</u>). After placing the animal at the start location, the experimenter quickly but calmly moved behind a black curtain next to the maze to not be visible to the animal during training trials.

Training consisted of two blocks: Build-Up and Updates. During probe sessions (each second session of a GL switch and additionally in Build-Up GL1: S6, GL2: S5, GL3-5 S4) there was no food in the maze for the first and ninth trial of the day and each time for the first 60 s of the trial to ensure that olfactory cues did not facilitate navigation to the GL. After 60 s food was placed in the GL while the animal was in a different part of the maze (to avoid the animal seeing the placement). All other trials of the day were run with food at the GL. Probe trials and GLs switches were initially minimized, to help shape the animal behaviour. In the first trial of the day, animals would not find food at the last presented location for both the first session of a new GL as well as probe trial days (e.g. always the second session of a new GL); thus these sessions were interleaved with normal training sessions with food present at the last known location in the first trial of the day to avoid the animals learning the rule that food is initially not provided.

To measure the animals’ performance, the actual path a mouse took was divided by the shortest possible path between a given start location and the GL, resulting in the log of normalized path length (Fig. 1B) and functioning as a score value. Given a sufficient food motivation and an established knowledge-network of the maze a mouse should navigate the maze efficiently. A score of 0 indicated that the mouse chose the shortest path and navigated directly to the goal. On average, animals would improve from a 3 times to 1.5-2 times longer path length than the shortest path, corresponding to 0.4 and 0.2-3 log values. Random walks through the maze are estimated with a model to result in a 4 times longer path (0.6 in log). The normalized path length of any first trial of a session was used to measure long-term memory since training sessions were two to three days apart.

First trial of the second sessions (probe trials) of each goal location in Build-up and Update phase were watched to score the number of times that animals crossed their current and previous goal location as well as the amount of time they dwelled there. As a control, same method was applied to two other nodes, one on the inner ring and the other on the outer ring of the maze. These nodes were selected in such a way that they were not close to each other and to the goal locations, with at least three gangways between them. Further, to control a false positive result, nodes that were in the way between goal locations were not chosen as a control Food motivation was ensured by restricting access to food for 12 h to 24 h before training and confirmed by both the number of trials ran each day as well as the count of trials during which the animal ate food at the first encounter with the food in each trial. If animals were not sufficiently motivated, the count of both would decrease. Additionally, animals were weighted three times a week and the average weekly weight was ensured to not fall below estimated 85% free-feeding weight, which was adapted for the normal growth of each animal across time.

### Behavior Analysis

The structure of the HexMaze experimental setup was reproduced as a directed graph with node numbering corresponding to the experimental one. Animal trajectories were thus described as sequences of visited nodes on this graph (Figure 1, top).

When measuring the distance of the animal location from one of the nodes in the optimal path, one has to consider the possible presence of multiple shortest paths of equal length connecting the start location with the goal. To take into account this source of ambiguity, for each trial, we computed the optimal path between the two locations using the HexMaze graph with weighted edges. The shortest path was computed multiple times each time on a different graph, first initialized with uniform weights and then adding small random noise to the value of every edge. In this way, in the presence of alternative and equivalent paths, the noisy weights would lead to the selection of either of the existing ones on a random basis. By collecting all the nodes happening to be described as belonging to a shortest path we thus obtain a list of all the nodes to be considered when computing the distance of the animal from the optimal path.

The experimental trials are divided as following: each condition (Build-Up, Location Update or Barrier Update) comprises 3 sessions. For each session we analyze separately the first trial and then the following trials in groups of 10 until trial number 31.

Distance from optimal path curves fitting:

We used the following function to parametrize the animal performance.

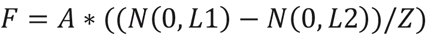

The difference of Gaussian functions (N) is normalized (Z) so that its maximum value is equal to. Therefore, A then controls the peak value of fit, while L1 and L2 its descending and ascending length, respectively. The fit is performed by optimizing the values of A, L1 and L2 (Figure 4, top).

### Simulations

The simulations are performed as following: we create a virtual HexMaze as a directed graph having the same structure of the real one. At every time step the virtual mouse moves from one node to an adjacent one. We do not allow trajectory reversals, so the node visited at the previous time step is not taken into account as a target. The start and goal locations are the same as those used in the experiment. Each run consists then in a sequence of nodes visited by the mouse and the run eventually ends when the animal reaches the goal. We augment the size of the modelled data by simulating multiple independent runs (n=50) for each experimental trial.

The movements of the virtual animal are generated according to an algorithm with two components: Random Search and the Foresight. The Random Search part consists in a procedure to select which node the animal is going to visit next and is meant to approximate an optimal search strategy. While performing Random Search the animal randomly picks the next node among the available ones. On top of this we introduce the possibility for the animal to take long diagonal runs that take it to another section of the maze. These diagonal runs are initiated with a probability η at any time step. If a diagonal run is initiated, then a node is randomly picked among those in the outer ring and at a distance of at least 3 steps from the current position of the animal. The mouse then uses the following time steps to reach this target along the shortest available path. Once the target is reached, the random node selection is resumed. We use different simulations to vary the value of η. Decreasing the value of this probability makes the search strategy approximate more and more a purely random walk through the environment. Higher values introduce a larger amount of “optimality” as they allow the animal to more quickly leave an already explored area.

The Foresight component on the other hand represents the ability of the animal to anticipate the location of the goal when getting within a certain distance from it. It is therefore aimed at representing the effect of experience and an increasing knowledge of the environment and of visual cues. At every step, we draw a random number from an exponential distribution with mean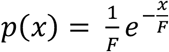. If the shortest path from the animal location to the goal node is smaller of this number, then the animal takes a direct path to the goal and the trial is over. Also in this case we run different sets of simulations varying the value of F. F=0 corresponds to an animal with no ability to remember the position of the goal from its current location, unless by running directly over it. As F increases the chances for the simulated mouse to detect the goal from some distance increase. Eventually a very large F would reproduce a goal-directed behavior.

For each set of parameters, we measure how well the statistics of the simulated runs reproduce those obtained from real animal behavior. To do so we use a combination of measures: 1) relative trial length (as the ratio between the actual length of the trajectory and the length of the shortest path between start and goal location); 2) maximal distance from the optimal path reached during the trial; 3) amount of time spent in the external ring of the maze vs. the internal ring. For each of these quantities we compare the distribution obtained from the experiment to the one generated simulating the trajectory of the animal according to a specific set of parameters. We measure the distance between the two distributions with Kolmogorov-Smirnov statistics. Therefore for each set of experimental trials we obtain get how well the statistical properties of the animal behavior can be reproduced by a certain choice of the model parameters.

The experimental trials are divided as following: for each condition (Build-Up, Location Update or Barrier Update) we used the first 3 session. During Build-Up more sessions were run for each goal location, however here we focus on the first 3 to be able to compare it with the Update phase. For each session we analyze separately the first trial and then the following trials in groups of 10 until trial number 31.

All analysis and simulations were performed using custom MATLAB code.

## Acknowledgments

Federico Stella is supported by the European Union’s Horizon 2020 research and innovation programme under grant agreement No. 840704 (BrownianReactivation), Alejandra Alonso by the European Union’s Horizon 2020 research and innovation programm under the Marie-Sklodowska-Curie grant M-Gate No. 765549.

